# Booster vaccination improves the durability of antibody-secreting plasma cells

**DOI:** 10.64898/2026.04.18.719387

**Authors:** An Qi Xu, Miu Shing Hung, Baizhi Chen, Miriam Llorian Sopena, Probir Chakravarty, Abdouramane Camara, Dinis Pedro Calado

**Affiliations:** Immunity and Cancer Laboratory, Francis Crick Institute, London, UK; Bioinformatics and Biostatistics, Francis Crick Institute, London, UK; West African Centre for Cell Biology of Infectious Pathogens, University of Ghana, Accra, Ghana; Immunity and Cancer Laboratory, The Babraham Institute, Babraham Research Campus, Cambridge, UK; Immunology Programme, The Babraham Institute, Babraham Research Campus, Cambridge, UK

## Abstract

Booster vaccination can restore antibody titres and protection, but whether it improves long-term durability by expanding plasma cell (PC) numbers or also by shifting PC fate toward intrinsically longer-lived states remains unclear. Here we established longitudinal in vivo ground truth for PC persistence by combining PC-specific genetic timestamping, clonal tracking, and multi-timepoint single-cell profiling across spleen and bone marrow. We resolved PC longevity as a layered, non-binary architecture comprising short-, intermediate-, and long-lived programs, and showed that program identity is specified early in secondary lymphoid tissues and largely maintained as PCs populate bone marrow niches. Primary vaccine responses initiated from naïve B-cells generated a prominent intermediate-lived wave, whereas memory B-cell recall during boosting redistributed output toward long-lived programs rather than recreating the intermediate-lived compartment characteristic of priming. Conserved longevity signatures projected onto early circulating PCs provide a cross-species framework to infer durability programs, supporting benchmarking of vaccine regimens by predicted persistence rather than peak titres.

**Highlights:**

- Genetic timestamping resolves short-, intermediate-, and long-lived PC programs
- Longevity programs are imprinted early and maintained from lymphoid organs to bone marrow
- Cross-species signatures stratify human blood and bone marrow PCs by persistence
- Boosting via MBC recall enriches long-lived PC and contracts the intermediate-lived tier

## Introduction

Public hesitancy toward booster vaccination is rising ^1–3^. Compared with primary vaccination, willingness to receive additional doses has fallen markedly, with >50% increases in hesitancy reported in some groups, driven by concerns about side effects, necessity and effectiveness ^1–3^. Inconsistent messaging and misinformation have further eroded confidence ^3,4^, prompting the World Health Organization to classify vaccine hesitancy as a major global health threat ^5^. Addressing this challenge requires mechanistic evidence at cellular and molecular resolution clarifying what boosters add beyond the first dose.

Durable humoral immunity rests on two cellular pillars: long-lived antibody-secreting plasma cells (PCs) and memory B-cells (MBCs). Primary vaccination engages naïve B-cells, generating both PCs and MBCs; booster doses predominantly re-engage pre-existing MBCs, which respond faster and differentiate into PCs more efficiently than naïve B cells ^6–10^. Although serology consistently shows that boosters increase antibody titres and protection ^9,11–15^, three long-standing questions remain unresolved. First, does PC persistence reflect a simple short-lived versus long-lived dichotomy, or discrete longevity programs? Second, when and where are these programs specified i.e., during PC differentiation in secondary lymphoid organs or later within bone marrow niches, and are they maintained across the tissue transition? Third, building on these points, does boosting act mainly by increasing PC numbers (quantity), or does it also re-shape the composition (quality) of PC persistence programs in a cell-of-origin dependent manner?

Progress has been limited because long-lived PCs lack a robust prospective definition in humans, and single time-point sampling cannot link transcriptional state to long-term survival outcomes across tissues ^16–22^. Here, we address these barriers by integrating PC-specific genetic timestamping and fate mapping with dense longitudinal single-cell transcriptomics and clonal tracking to link transcriptional state and clonal history to durability across spleen and bone marrow. We define a layered, non-binary architecture of PC persistence programs, show that these programs are imprinted early and largely maintained across tissues, and provide conserved cross-species signatures that stratify human PCs in bone marrow and blood. Finally, by combining human vaccination datasets with mouse adoptive-transfer models that isolate MBC-derived recall, we show that boosting redistributes PC output toward intrinsically long-lived programs while selectively contracting an intermediate-lived compartment characteristic of priming.

## Results

### A Layered Architecture of PC Longevity

To define PC age in vivo, we used a PC-specific genetic fate-mapping system (*Jchain^creERT2^* mice) in which Cre recombinase is driven by the *Jchain* promoter and reported by GFP expression ^23^. Tamoxifen administration activates Cre and irreversibly induces RFP ^24^, thereby selectively and permanently labeling PCs generated within the dosing window ^23^ (**Figure 1A**).

**Figure 1.**
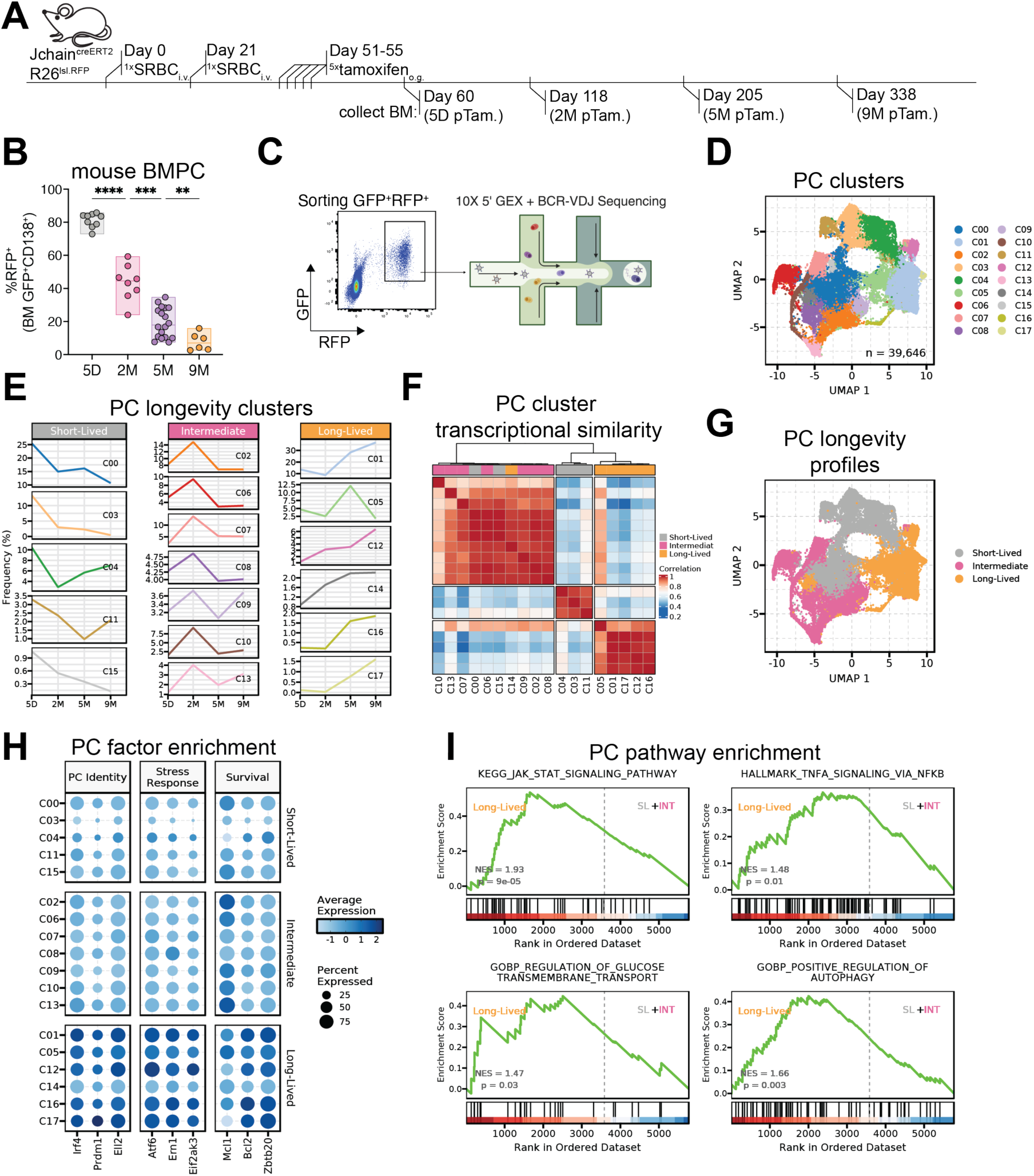
A Layered Architecture of Plasma Cell Longevity. **(A)** Schematic of the experimental design. **(B)** Cumulative frequency of RFP⁺ timestamped cells among GFP⁺ plasma cells in the bone marrow (BM). **(C)** Gating strategy for sorting GFP⁺RFP⁺ timestamped bone marrow plasma cells (BMPCs) and downstream workflow for single-cell gene-expression profiling and paired B-cell receptor (BCR) VDJ sequencing. **(D)** UMAP projection of BMPCs from all-time points, clustered into 18 transcriptional states (C0-C17). **(E)** Classification of BMPC clusters into long-lived, intermediate or short-lived program, based on the abundance of cells bearing each longevity signature across the 9-month time frame. **(F)** Jaccard index–based clustering of BMPC states according to transcriptional similarity. **(G)** UMAP of BMPCs colored by longevity program. **(H)** Relative mRNA expression of representative genes associated with plasma cell identity and function across states, grouped by longevity program. **(I)** Gene set enrichment analysis (GSEA) of pathways implicated in plasma cell survival, comparing cells with long-lived program to those with short-lived and intermediate-lived program. Statistical analysis in (B) was performed using Brown-Forsythe and Welch one-way ANOVA with Dunnett’s T3 multiple comparisons test; ****P < 0.0001, **P < 0.01, *P < 0.05. GSEA in (I) reports normalized enrichment scores (NES) and P values.

To build a longitudinal atlas of PC transcriptional states, we profiled PCs generated during polyclonal primary and booster responses to a T-dependent antigen (sheep red blood cells, SRBC). We traced timestamped RFP⁺ PCs in the bone marrow at four time points after induction (5 days, 2 months, 5 months, and 9 months). As expected, >80% of bone marrow PCs were RFP⁺ at day 5, reflecting recent labeling, whereas <20% remained RFP⁺ at 9 months, consistent with progressive attrition and selective persistence of longer-lived cells (**Figure 1B**). Given the relatively slow turnover of bone marrow PCs (BMPCs) compared with peripheral compartments ^21–23,25,26^, this niche provides a stable setting to follow survival trajectories over a substantial fraction of the murine lifespan.

We FACS-sorted RFP⁺ BMPCs as GFP⁺RFP⁺ for 10x Genomics single-cell RNA-seq and paired B-cell receptor (BCR)-VDJ sequencing (**Figure 1C**). After quality control and integration, we obtained 39,646 BMPCs across all time points. Unsupervised clustering based on transcriptional similarity resolved 18 transcriptional states (**Figure 1D**; **Figure S1A**). To organize these states by longevity, we quantified how state abundance changed over time (**Figure 1E**). We next defined longevity programs as a set of transcriptional states that share a reproducible temporal abundance trajectory across months. Silhouette scoring and hierarchical clustering (**Figure S1B, C)** identified three recurrent temporal trajectories that define three longevity programs: short-, intermediate-, and long-lived. Short-lived states (C0, C3, C4, C11, C15) dominated at day 5 and were largely absent by 2 months. Intermediate-lived states (C2, C6, C7, C8, C9, C10, C13) peaked at ∼2 months and declined by 5 months. Long-lived states (C1, C5, C12, C14, C16, C17) were relatively infrequent early but were enriched at 5-9 months. Together, these dynamics reveal a layered, non-binary architecture of PC longevity rather than a simple short- versus long-lived dichotomy.

Consistent with this organization, unsupervised similarity heatmap showed that states within each longevity program are more similar to one another than to states in other programs (**Figure 1F**), and UMAP projection revealed clear transcriptional separation between the three longevity programs (**Figure 1G**). Thus, BMPCs partition into discrete transcriptional states with distinct survival trajectories, including a prominent intermediate-lived compartment not anticipated by classical models, an architecture that becomes apparent only through multi-timepoint fate mapping.

### Conserved Survival Modules in Long-Lived PCs

To identify molecular features associated with persistence, we inspected state-defining genes across the three longevity programs. Long-lived PCs were characterized by a coordinated survival module that reinforced PC identity, buffered secretory stress, and engaged pro-survival circuitry. Core PC identity regulators (*Irf4*, *Prdm1*) ^27–30^ and the secretory factor *Ell2* ^31^ were most strongly expressed in states in the long-lived program (**Figure 1H**). These same states upregulated key unfolded protein response components (*Atf6*, *Ern1*, *Eif2ak3*) ^32–35^, consistent with increased capacity to manage endoplasmic-reticulum stress. Respecting anti-apoptotic factors, *Mcl1* ^36^, was elevated in intermediate- and long-lived states, whereas *Bcl2* and *Zbtb20* ^36–38^ were selectively enriched in long-lived states (**Figure 1H**). At the pathway level, gene set enrichment analysis showed that long-lived states were enriched for signatures linked to PC maintenance, including JAK/STAT and NF-κB signaling ^39,40^, glucose transport ^41^, and autophagy ^17,42,43^ (**Figure 1I**). Together, these features define a conserved, persistence-associated module in which reinforced PC identity and secretory capacity are coupled to UPR-mediated stress buffering and pro-survival signaling; PCs that most strongly engage this module are those that preferentially occupy long-lived states.

Collectively, these data place PC transcriptional states within a layered longevity architecture comprising short-, intermediate-, and long-lived programs that are inferred from their abundance dynamics over time. By integrating long-term in vivo fate mapping with single-cell transcriptomics, we provide a temporal atlas of PC states and a mechanistic framework in which PC lifespan tracks with engagement of a conserved survival module, offering a basis for benchmarking and ultimately modulating durable humoral immunity.

### Cross-Species Longevity Signatures Stratify Human PCs

Genetic fate mapping enables direct measurement of PC persistence in mice, but equivalent temporal labeling is not feasible in humans, where long-lived PCs have instead been inferred from surface phenotype. Human CD38^hi^ BMPCs can be partitioned into three subsets (“A”: CD19^+^CD138^−^, “B”: CD19^+^CD138^+^, “D”: CD19^−^CD138^+^) ^16–18^. In bone marrow aspirates from healthy adult donors (24-55 years), populations A and B varied in size, whereas population D was consistently small but detectable across individuals (**Figure S2A, B**). CD19 downregulation has been proposed as a correlate of extended PC survival ^16,17,22,44,45^ and several studies reported enrichment of long-lived, antigen-specific PCs in population D ^16–18^. However, antigen-specific PCs (for example, vaccinia-specific cells) are also found in population B ^20^, indicating that CD19 loss is associated with, but does not uniquely define, long-lived PCs ^17,18^, leaving the surface phenotype of human long-lived PCs unresolved.

To overcome this limitation, we projected mouse-derived longevity signatures onto a single-cell RNA-seq dataset of human BMPCs from healthy donors. All three longevity programs, short-, intermediate-, and long-lived, were readily identifiable within the human BMPC compartment (**Figure 2A**), indicating that the underlying transcriptional architecture is conserved across species. Human PCs assigned to the long-lived program showed elevated expression of core PC identity regulators (*IRF4*, *PRDM1*), secretory machinery (*ELL2*), unfolded protein response components (*ATF6*, *ERN1*, *EIF2AK3*), and anti-apoptotic factors (*MCL1*, *BCL2*, *ZBTB20*) (**Figure 2B**). They were also enriched for the same maintenance pathways as murine long-lived PCs, including JAK/STAT and NF-κB signaling, glucose transport, and autophagy (**Figure 2C**). This concordance supports mouse longevity signatures as molecular proxies for stratifying human PCs by predicted persistence.

**Figure 2.**
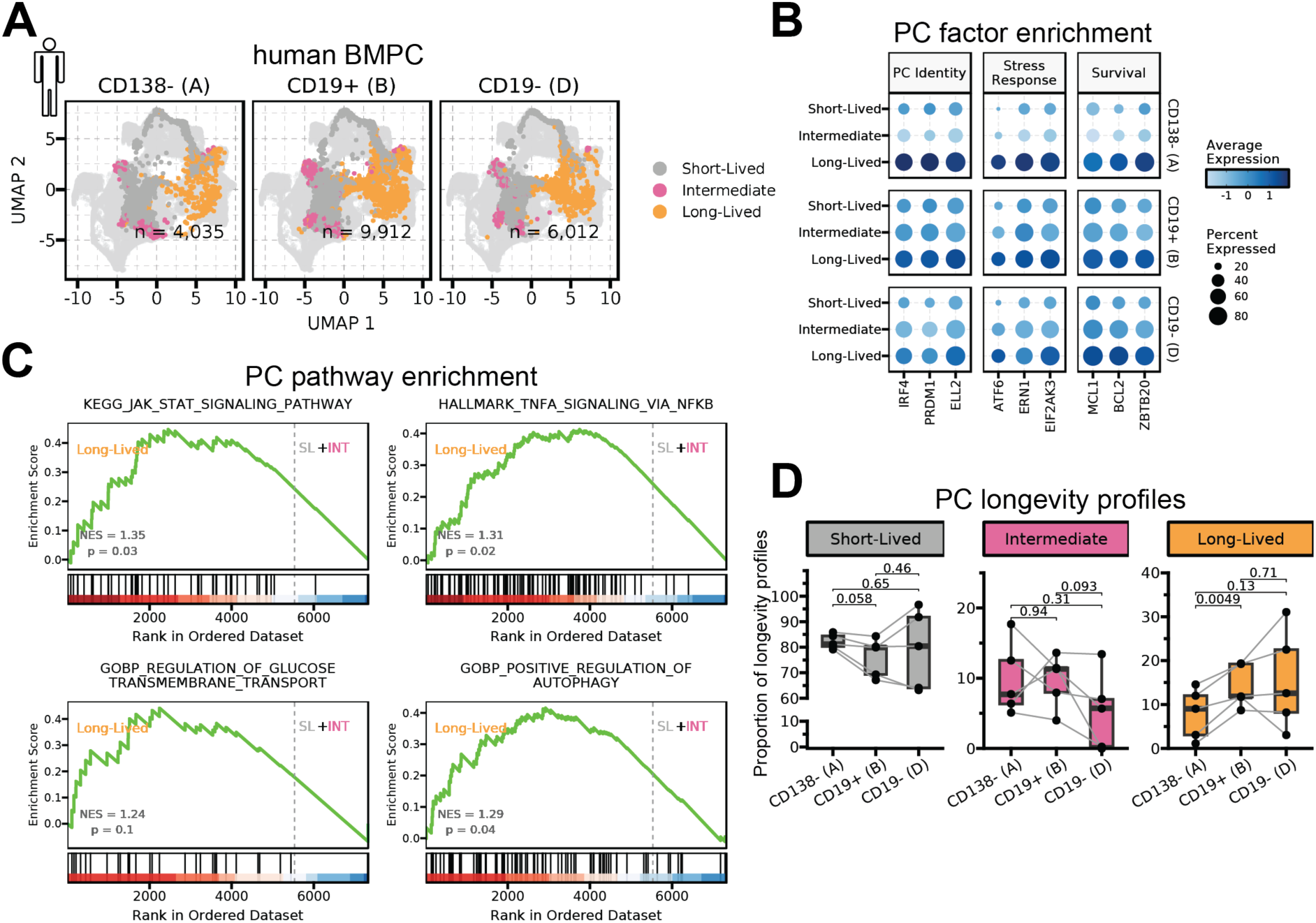
Cross-Species Longevity Signatures Stratify Human PCs. **(A)** Reference mapping of CD38⁺ human bone marrow plasma cell (BMPC) populations A (CD138⁻CD19⁺), B (CD19⁺CD138⁺) and D (CD19⁻CD138⁺) onto murine BMPC states defined in Figure 1. **(B)** Relative mRNA expression of genes associated with plasma cell identity and function in the three human BMPC populations, stratified by inferred longevity program. **(C)** Gene set enrichment analysis (GSEA) of pathways implicated in plasma cell survival, comparing human BMPCs mapped to long-lived program with those mapped to short-lived and intermediate programs. **(D)** Comparison of long-lived, intermediate-lived and short-lived program abundances between populations A, B and D using paired t-tests (P values shown).

Having established conservation, we asked how longevity programs distribute across the A/B/D subsets. Although most human BMPCs mapped to the short-lived program, populations B and D contained a higher fraction of long-lived PCs (**Figure 2D**), consistent with prior reports ^16–18,20^. Together, these data support a conserved, signature-based cross-species framework for stratifying BMPCs by predicted lifespan even in the absence of definitive surface markers.

### Early Imprinting of PC Longevity

A central question is whether transcriptionally defined longevity programs arise only after PCs enter bone marrow niches or are already specified in the secondary lymphoid organs where PCs are generated. To address this, we analyzed an independent single-cell RNA-seq and BCR-VDJ dataset profiling spleen and bone marrow PCs ^21^ and projected these cells onto our atlas of short-, intermediate-, and long-lived states. Seventeen of our 18 transcriptional states were recovered in the independent dataset (**Figure 3A**), indicating robust reproducibility of the atlas. The few differences were limited to rare states consistent with sampling depth (C16/C17).

**Figure 3.**
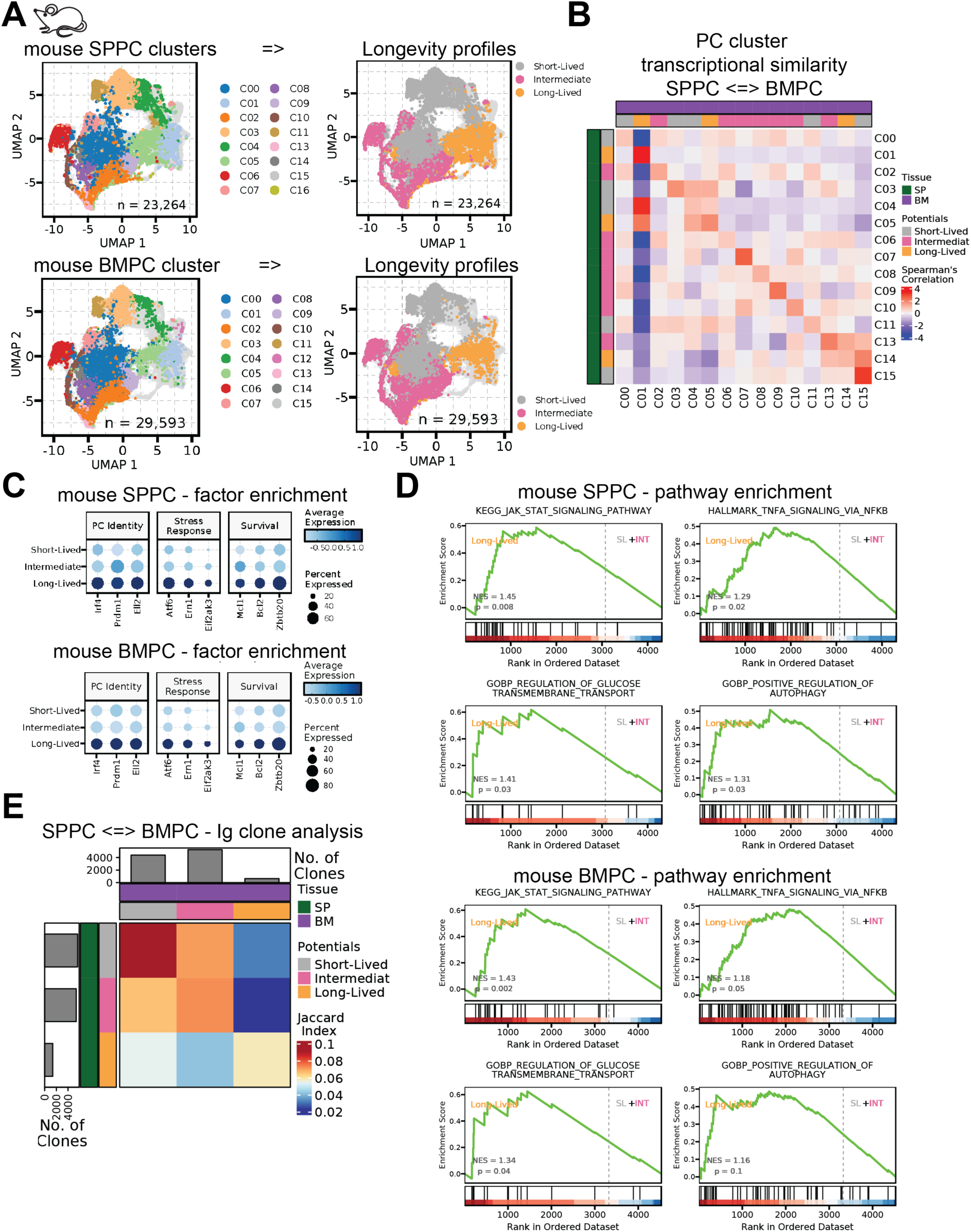
Early Imprinting of Plasma Cell Longevity. **(A)** UMAP projections of reference-mapped splenic plasma cells (SPPCs; top) and bone marrow plasma cells (BMPCs; bottom), colored by transcriptional cluster identity (left) or inferred longevity program (right). **(B)** Transcriptional similarity between reference-mapped SPPC and BMPC clusters, quantified by Spearman’s correlation. **(C)** Relative mRNA expression of representative signature genes in reference-mapped SPPCs (top) and BMPCs (bottom), grouped by longevity program. **(D)** Gene set enrichment analysis (GSEA) of pathways associated with plasma cell survival in reference-mapped SPPCs (top) and BMPCs (bottom), comparing cells mapped to long-lived program with those mapped to short-lived or intermediate programs. **(E)** BCR-VDJ analysis identifying convergent clones, defined by shared heavy-chain V and J gene usage and identical CDR3 sequence, occurring within the same longevity program in spleen and bone marrow.

With this overall concordance, we next asked whether state identity is preserved as PCs transit from spleen to bone marrow. Transcriptional similarity analysis showed that each splenic state was most closely related to its namesake bone marrow state (**Figure 3B**), indicating preservation of cluster-level gene-expression signatures as PCs transit between tissues. Consistent with this, splenic and bone marrow PCs mapped to the long-lived program expressed high levels of PC identity and unfolded protein response genes (*Ell2, Irf4, Prdm1, Atf6, Ern1, Eif2ak3*), together with pro-survival factors (*Mcl1, Bcl2, Zbtb20*) (**Figure 3C**). Pathway analysis likewise showed convergent enrichment of JAK/STAT, NF-κB, glucose-transport, and autophagy pathways in long-lived PCs from both tissues (**Figure 3D**), indicating engagement of the same conserved survival module independent of anatomical site. Clonal analysis further supported early specification of longevity: clones shared between spleen and bone marrow overwhelmingly occupied the same longevity program rather than redistributing across programs (**Figure 3E**). If bone marrow niches routinely imposed longevity de novo, individual clones would be expected to split across different longevity programs when comparing the two tissues.

Together, these findings indicate that longevity programs are already imprinted in secondary lymphoid organs and are largely maintained as PCs enter and persist within bone marrow niches. While niche-derived signals may modulate survival, the dominant features of longevity programs appear to be specified early during PC differentiation and conserved across tissues.

### Boosters Reshape the PC Longevity Landscape

Because longevity programs are broadly conserved across tissues, circulating PCs offer a practical window into durability programs, enabling longitudinally anchored interpretation of early blood PC states after vaccination. We therefore analyzed single-cell transcriptomic data of circulating PCs from SARS-CoV-2–naïve individuals receiving mRNA-1273, sampling blood at day 14 after dose 1 and day 6 after dose 2 (Figure. 4A). Integrated analysis of circulating PCs identified all three longevity programs: short-, intermediate-, and long-lived, at both time points (**Figure 4B**).

**Figure 4.**
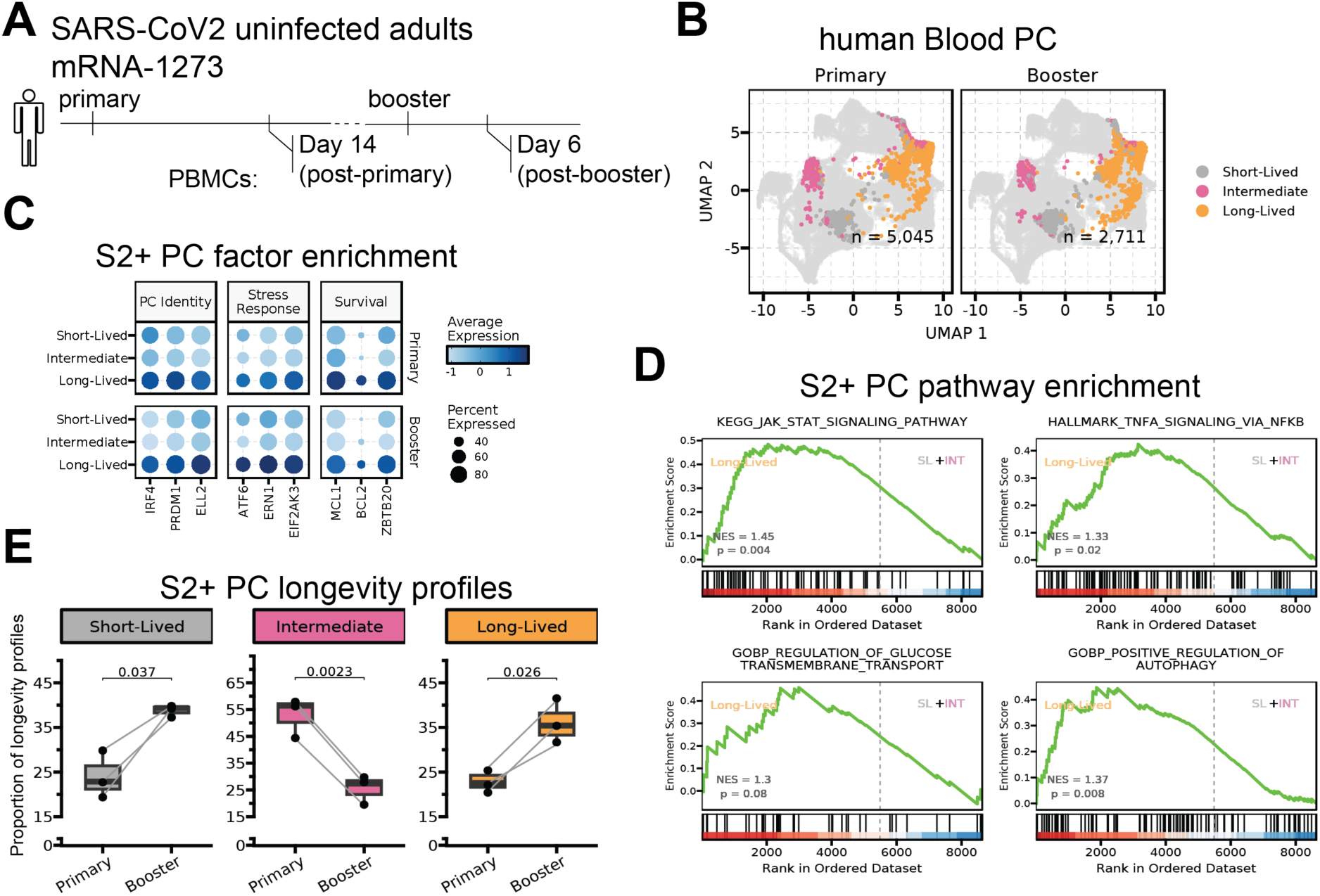
Boosters Reshape the Plasma Cell Longevity Landscape. **(A)** Schematic of sample composition and sampling schedule, showing blood collection at day 14 after the first (primary) vaccine dose and day 6 after the second (booster) dose. **(B)** UMAP projection of reference-mapped plasma cells (PCs) from primary and booster responses, colored by inferred longevity program **(C)** Relative expression of representative signature genes associated with PC identity and function in S2⁺ PCs, grouped by longevity program and immunization phase (primary vs booster). **(D)** Gene set enrichment analysis (GSEA) of pathways associated with PC survival in S2⁺ PCs, comparing cells mapped to long-lived program with those mapped to short-lived (SL) or intermediate (IN) programs. **(E)** Composition of S2⁺ PCs by transcriptional cluster and longevity program in primary and booster responses, with statistical comparison of longevity program frequencies between primary and booster samples using paired t-tests (P values shown).

To focus on vaccine-induced cells, we identified Spike-specific (S2^+^) PCs (**Figure S3**). S2^+^ PCs assigned to the long-lived program expressed the same hallmark genes (**Figure 4C**) and survival pathways (**Figure 4D**) that defined cross-species long-lived PCs (**Figure 1H, I**; **Figure 2B, C**; **Figure 3C, D**), supporting the applicability of the longevity signatures to circulating human PCs.

Direct comparison of primary and booster vaccination showed that boosting changes the composition of the S2⁺ PC pool. Although priming is often assumed to produce a largely short-lived burst, circulating S2⁺ PCs after primary vaccination were instead dominated by the intermediate-lived program. After boosting, long-lived S2⁺ PCs increased sharply, short-lived PCs also rose, and the intermediate-lived compartment contracted (**Figure 4E**), indicating a qualitative redistribution of longevity programs.

Together, these data show that intermediate-lived PCs are a defining feature of the primary response and are selectively reduced after boosting, which instead favors PCs committed to short-and, critically, long-lived programs. Booster vaccination therefore reshapes the distribution of PC longevity programs in humans, providing a cellular basis for enhanced protection durability after boosting and further supporting a layered architecture of PC longevity.

### MBC Recall Preferentially Generates Long-Lived PCs

The redistribution of PC longevity after boosting likely reflects the distinct B-cell inputs engaged during priming versus recall. To test how MBCs contribute to PCs with different longevity programs, we used an adoptive-transfer system based on B1-8^hi^ B-cells, which express a high-affinity BCR specific for 4-hydroxy-3-nitrophenylacetyl (NP) ^47^ (**Figure 5A**). This design enables direct comparison of PCs generated during a primary response initiated from naïve B-cells with PCs generated from antigen-experienced MBCs upon recall to the same antigen.

**Figure 5.**
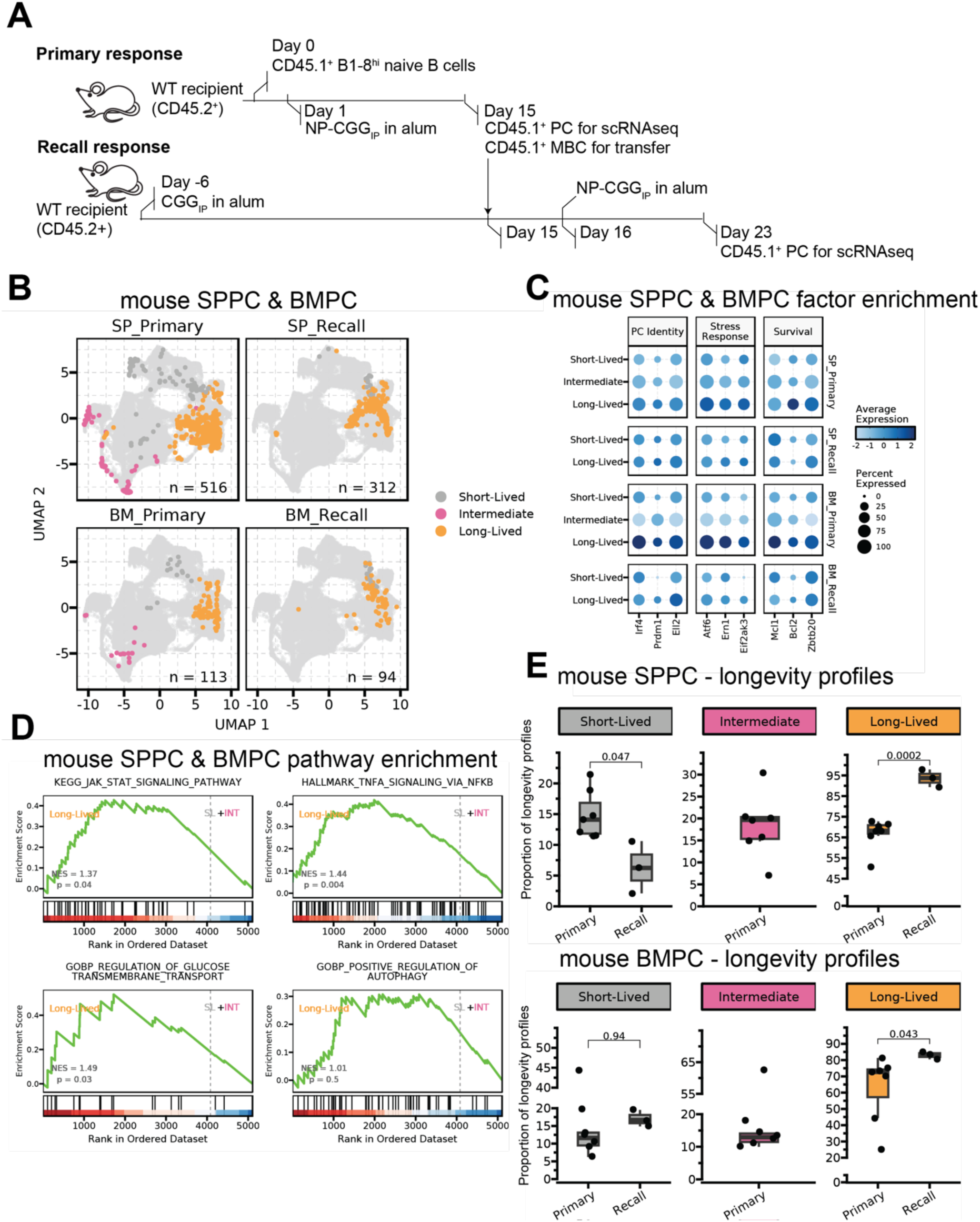
MBC Recall Preferentially Generates Long-lived Plasma Cells. **(A)** Schematic of the immunization and sampling strategy. **(B)** UMAP projection of reference-mapped plasma cells from primary and recall responses in spleen and bone marrow, colored by longevity program. **(C)** Relative expression of representative signature genes associated with plasma cell identity and function across longevity programs. **(D)** Gene set enrichment analysis (GSEA) of pathways associated with plasma cell survival, comparing cells mapped to long-lived program with those mapped to short-lived or intermediate programs. **(E)** Comparison of the proportions of plasma cell clusters and longevity programs between primary and MBC-recall responses in SP (top) and BM (bottom), analyzed using unpaired Student’s t-tests (P values shown).

To model primary vaccination, CD45.1^+^ B1-8^hi^ naïve B-cells were transferred into wild-type CD45.2^+^ recipients and immunized with NP-chicken gamma globulin (NP-CGG) in alum (**Figure 5A**). Fourteen days later, we detected robust donor-derived NP-specific responses (**Figure 5A**; **Figure S4A**), and FACS-purified donor-derived PCs from spleen and bone marrow for single-cell RNA-seq. In parallel, we isolated donor-derived MBCs from the same primary recipients and transferred them into a second cohort of antigen-primed CD45.2^+^ mice (**Figure 5A**). These mice were boosted with NP-CGG, and seven days later we sorted MBC-derived PCs from spleen and bone marrow for single-cell RNA-seq (**Figure 5A**; **Figure S4A**). Thus, differences between primary and recall outputs reflect the intrinsic output of naïve-driven priming versus MBC-driven recall under matched antigenic conditions.

We indexed PCs by response type (primary versus MBC-derived recall), tissue, and mouse using hashtag labeling. Integrated transcriptomic analysis identified all three longevity programs: short-, intermediate-, and long-lived, across conditions (**Figure 5B**; **Figure S4B**). As in our atlas and human projections, PCs assigned to the long-lived program expressed conserved hallmark genes (**Figure 5C**) and enriched survival pathways (**Figure 5D**). We next compared fate allocation between priming and recall. In spleen, MBC-derived recall responses were strongly biased toward long-lived PCs, with only a small minority adopting a short-lived program and no cells acquiring an intermediate-lived profile (**Figure 5E**). In contrast, primary responses produced a mixed output spanning short-, intermediate-, and long-lived programs. Notably, the absence of intermediate-lived PCs from the MBC-derived response mirrors the contraction of this compartment after booster vaccination in humans (**Figure 4E**). These early differences in fate within secondary lymphoid tissue propagated to the bone marrow (**Figure 5E**). Naïve-driven primary responses seeded the bone marrow with a broad mixture of short-, intermediate-, and long-lived PCs, whereas MBC-driven recall responses generated a BMPC pool enriched for long-lived PCs and depleted of intermediate-lived cells (**Figure 5E**).

Together with the human vaccination data (**Figure 4**), these experiments indicate that MBC recall during boosting preferentially yields long-lived PC output while disfavoring intermediate-lived PC formation, with immediate consequences for bone marrow composition.

## Discussion

Long-lived PCs are essential for durable antibody mediated protection. How their lifespan is specified, and how booster vaccination improves durability, has remained unresolved. By combining PC-specific genetic timestamping with longitudinal single-cell transcriptomics and clonal tracking, and by projecting conserved signatures onto human bone marrow and blood, we resolve PC persistence into a layered, non-binary architecture and identify MBC recall as a privileged route to long-lived PC fate.

A central finding is that PC longevity does not reduce to a short-lived versus long-lived dichotomy. Time-resolved fate mapping resolves discrete transcriptional states that group into three longevity programs: short-, intermediate-, and long-lived; revealing a prominent intermediate-lived tier that persists for months but does not reach the longest-lived state. This intermediate tier emerges from unbiased clustering, is present in both spleen and bone marrow, and is detectable in human bone marrow and blood. Resolving this intermediate-lived compartment refines the long-standing “short versus long” model and explains how priming can sustain antibody titres for months without immediately committing most output to the longest-lived fate. A layered architecture may therefore provide functional flexibility: intermediate-lived PCs could maintain titres while long-lived PCs are selected and consolidated, preserving finite niche capacity for higher-fitness PCs generated by subsequent exposures.

Our data further indicate that longevity programs are specified early and largely maintained across tissues. Splenic PCs map to cognate bone marrow states, and shared clones overwhelmingly occupy the same longevity program in both sites. This argues against a model in which bone marrow niches routinely reprogram newly arrived PCs into long-lived cells de novo. Instead, longevity appears to be imprinted during or shortly after PC differentiation in secondary lymphoid organs and carried into the bone marrow. This does not exclude a modulatory role for niche-derived cues; rather, it suggests they act primarily as permissive, sustaining inputs that enable persistence of pre-specified programs, refining, rather than establishing, PC longevity.

Across tissues and species, long-lived PCs converge on a conserved survival module that integrates reinforced PC identity and secretory capacity with stress buffering and pro-survival circuitry. In spleen, blood and bone marrow, and in mouse and human, long-lived PCs upregulate core PC regulators and secretory machinery, engage unfolded protein response components, and enrich survival-associated pathways alongside anti-apoptotic factors. This architecture is consistent with how long-lived PCs sustain high-rate antibody secretion without succumbing to chronic proteotoxic stress, and it provides a conserved molecular framework to stratify PCs by predicted persistence across species.

Leveraging this framework, we reveal that booster vaccination improves PC quality, not simply a larger response. In vaccinated individuals, circulating PCs after priming were unexpectedly enriched for the intermediate-lived program, contrary to the prevailing view that the primary blood response is dominated by a transient, short-lived burst. After boosting, the distribution shifted sharply: long-lived programs expanded while the intermediate-lived compartment contracted. Mirroring this in a controlled setting, mouse adoptive-transfer experiments that isolate antecedent B-cell origin showed that naïve-driven primary responses generate short-, intermediate-, and long-lived PC programs, whereas MBC-driven recall during boosting is strongly biased toward long-lived PCs: not forming the intermediate-lived wave characteristic of priming. Together, these concordant human and mouse findings resolve a long-standing question: boosters do not merely increase the quantity of antibody-secreting PCs, they improve PC quality by biasing fate toward longer-lived programs, and they do so through preferential through MBC recall.

These findings have practical implications for vaccine evaluation and design. By linking transcriptional state and clonal history to survival outcomes over months, supported by fate mapping and clonal continuity across tissues, this work moves beyond descriptive PC heterogeneity to a predictive framework for persistence. First, the longevity signatures provide cross-species biomarkers that can be projected onto human blood to stratify vaccine-induced PCs by predicted lifespan from early time points, supporting benchmarking of vaccine platforms, adjuvants and booster schedules by durability programs rather than peak antibody titre. Second, because longevity programs are conserved across tissues, early circulating PCs may offer a minimally invasive readout of eventual bone marrow composition, providing a prospective surrogate endpoint for durability. This is particularly valuable for optimizing strategies in populations at risk of waning immunity, including older adults, immunodeficient individuals and patients receiving B-cell depleting therapies. More broadly, this framework should extend to settings where durable antibody production is beneficial or harmful, including therapeutic vaccination, chronic infection, autoimmunity and cancer, where selectively expanding or limiting long-lived PC output could have sustained clinical impact.

## Acknowledgements

We thank the members of the Immunity and Cancer laboratory [FCI, London, UK], Amalie Grenov, Carola Vinuesa [FCI, London, UK], Martin Turner [Babraham Institute, Cambridge, UK] for critical discussions and comments. We thank the FCI scientific platforms (Biological Resource Facility, Flow Cytometry, Advanced Sequencing) for expert advice and technical support. Schematics were created with BioRender.

## Author contributions

Conceptualization: A.Q.X., M.S.H., D.P.C.

Methodology: A.Q.X., M.S.H., B.C., M.L.S., P.C., A.C.

Investigation: A.Q.X., M.S.H., B.C., M.L.S., P.C., A.C.

Resources: D.P.C.

Visualization: A.Q.X., M.S.H., D.P.C.

Funding acquisition: D.P.C., A.Q.X., A.C.

Supervision: D.P.C.

Writing-original draft: A.Q.X., M.S.H., D.P.C.

Writing-review and editing: All authors.

## Data, code, and materials availability

All data are available in the main text or the supplementary materials.

## Supplemental information

### Materials and Methods

#### Mice

All mice in this study were bred and maintained on the C57BL/6 background at the Francis Crick Institute biological research facility under specific pathogen-free conditions. The *Jchain*^creERT2^ mouse strain is previously described^23^ and was crossed to carry a *R26*^lsl.RFP^ allele^24^. For PC timestamping experiments, *Jchain*^creERT2^ *R26*^lsl.RFP^ female and male mice were used. For cell transfer experiments, mice heterozygous for the *Igh*^B1–8hi^ allele^47^ were crossed to carry the CD45.1 congenic allele. *Igh*^B1–8hi^ donor female and male mice were used and sex-matched wildtype recipients. All animal experiments were conducted in compliance with national and institutional animal care guidelines and received approval from the Strategic Oversight Committee of the Biological Research Facility at The Francis Crick Institute, which includes the Animal Welfare and Ethical Review Body, as well as the UK Home Office. All animal care and procedures adhered to the UK Home Office regulations outlined in the Animals (Scientific Procedures) Act 1986 and were approved by the Biological Research Facility at The Francis Crick Institute.

#### Human sample acquisition

Human bone marrow mononuclear cells (BM MNCs) were acquired frozen from STEMCELL Technologies (Catalog #70001). All samples were obtained under informed consent and protocols approved by an Institutional Review Board, the U.S. Food and Drug Administration, the U.S. Department of Health and Human Services, or an equivalent regulatory authority. The use of these samples in this study was approved by the Human Biology Facility at The Francis Crick Institute, which operates under the Human Tissue Authority in accordance with the Human Tissue Act. BM MNCs were derived from eight adult donors with the following characteristics: ages ranged from 24 to 55 years; four donors were female and four were male. Donors self-identified as Caucasian (n = 3), African American (n = 2), Hispanic (n = 1), or mixed ethnicity (n = 2). Individual donor details, including lot number, age, sex, and ethnicity, are listed as follows: 2404410012 (46, male, Caucasian), 2408416007 (55, female, Caucasian), 2405402006 (48, female, Caucasian), 2412402002 (24, female, African American), 2412402003 (37, male, mixed), 2411406008 (54, female, Hispanic), 2412402009 (52, male, African American), and 2412404004 (54, male, mixed).

#### Flow cytometry and cell sorting

Freshly isolated or thawed cells were kept as single cell suspension in ice cold FACS buffer prior to staining. Cells were stained in 10 ul antibody cocktail per 1 million cells. Fc block was applied first before any staining. For intracellular staining, cells were first stained with surface antibodies and then fixed with 4% formaldehyde (PFA, Thermo Scientific, Cat#: 28908) for 15 minutes at room temperature, washed once with PBS, then fixed with True-NuclearTM Transcription Factor Buffer Set (BioLegend, Cat# 424401) for 20 minutes at 4°C and kept in True-Nuclear 1X Perm Buffer for downstream intracellular staining. Flow cytometry data was acquired on BD FACSymphony A5, BD LSRFortessa X-20 using FACS-Diva software (BD Biosciences), or on Sony ID7000 using its system software. Sorting was performed using BD FACSAria Fusion Flow Cytometer and data acquired using FACS-Diva software (BD Biosciences). Hashtagged cells were sorted at 4°C and kept in PBS + 0.04% BSA for downstream scRNAseq analysis, or in RPMI (Gibco, cat# 61870) for adoptive cell transfer. All flow cytometry data were analyzed on FlowJo (BD, v10.10.0).

#### Antibodies and reagents

For flow cytometry and cell depletion experiments, the following antibodies and reagents were used. Human cells were stained with antibodies against CD138 (MI15, BioLegend, 356554, PE/Fire700; 5 µl per test), CD14 (M5E2, BioLegend, 301826, biotin; 1:200), CD19 (SJ25C1, BioLegend, 363028, BV785; 1:200), CD27 (L128, BD Biosciences, 563815, BUV395; 1:200), CD3 (UCHT1, eBioscience, 13-0038-82, biotin; 1:200), CD38 (HIT2, BD Biosciences, 612947, BUV496; BioLegend, 303532, BV605; 1:200), and IgD (IA6-2, BioLegend, 348220, BV510; 1:200). Human Fc receptors were blocked using Human Fc Block (Fc1, BD Biosciences, 564219; 1:200). Mouse cells were stained with antibodies against B220 (RA3-6B2, BD Biosciences, 612838, BUV737; BioLegend, 103248, BV510; 1:200), CD138 (281- 2, BD Biosciences, 740240, BUV395 and 569692, BV786; BioLegend, 142506, APC; 1:200), CD19 (6D5, BioLegend, 115528, AF700 and 115506, FITC; 1:200; or 1D3, BD Biosciences, 563557, BUV395; 1:200), CD38 (90, BD Biosciences, 740361, BV605; 1:200), CD45.1 (A20, BioLegend, 110732, BV421; 1:100), CD95 (Jo2, BD Biosciences, 740906, BV786; 1:200), IgD (11-26c.2a, BioLegend, 405731, BV711; 1:200), IgG1 (A85-1, BD Biosciences, 560089, APC and 741733, BUV737; 1:200), and IgM (II/41, eBioscience, 25-5790, PE-Cy7; 1:300). Fc receptors were blocked using anti-mouse CD16/32 (2.4G2, BD Biosciences, 553141; 1:200). For in vivo or ex vivo depletion experiments, biotin-conjugated antibodies against CD11c (N418, BioLegend, 117304; 1:1000), CD4 (GK1.5, BioLegend, 100404; 1:1000), CD45.2 (104, BioLegend, 109804; 1:500), CD8a (53-6.7, BD Biosciences, 553029; 1:1000), GL7 (GL-7, BioLegend, 144616; 1:1000), IgD (11-26, eBioscience, 13-5993-85; 1:1000), NK1.1 (PK136, BioLegend, 108704; 1:1000), and Ter119 (TER-119, BioLegend, 116204; 1:1000) were used. NP-specific B cells were detected using NP-PE (ChemCruz, sc-396483; 1:500). Biotinylated antibodies were revealed using streptavidin-AF700 (Invitrogen, S21383; 1:1000) or streptavidin-BUV737 (BD Biosciences, 612775; 1:200). Sample multiplexing was performed using TotalSeq™ anti-mouse hashtag antibodies 1–9 and 11 (M1/42 and 30-F11, BioLegend, 155861–155877 and 155881; 1:500). Cell viability was assessed using Zombie NIR™ Fixable Viability Dye (BioLegend, 423106; 1:200).

#### Immunization, tamoxifen administration and cell adoptive transfers

For SRBC immunization, *Jchain*^creERT2^ *R26*^lsl.RFP^ mice were injected with 1 x 10^9^ defibrinated sheep red blood cells (SRBCs, TCS Biosciences Ltd, Cat# SB054) in phosphate-buffered saline (PBS, Gibco, Cat# 11503387) intravenously. To induce plasma cell timestamping, 4 mg tamoxifen (Sigma-Aldrich, Cat# T5648) dissolved in sunflower seed oil (Cat# S5007-250mL, Sigma-Aldrich) were administered by oral gavage once per day for five days. To generate mouse BMPC longevity signatures, timestamped PCs (GFP^+^RFP^+^) were sorted from BM harvested at 5 days, 2 months, 5 months and 9 months post tamoxifen treatment and subjected to scRNA-seq.

For *Igh*^B1–8hi^ B-cell adoptive transfer experiments, naïve B-cells were isolated from spleens of *Igh*^B1–8hi^ CD45.1^+^ mice using CD43 MicroBeads (Miltenyi Biotec, Cat#130-049-801). The percentage of NP-reactive B-cells were determined using flow cytometry and 2 million NP-reactive naïve B-cells were transferred into wildtype CD45.2^+^ sex-matched recipients. One day after cell transfer, *Igh*^B1–8hi^ B-cell recipient mice were injected intraperitoneally with 100 ug of alum-precipitated NP_15_-CGG (Biosearch Technologies, N-5055B) in 100 ul of PBS. Splenic and BM cells (femur and tibia) were isolated 14 d.p.i and enriched for donor-derived cells by depleting CD45.2^+^ recipient cells, T-cells, NK cells, naïve B-cells, GC B-cells, myeloid cells and erythrocytes using a mix of biotinylated anti-CD45.2, CD4, CD8, NK1.1, IgD, GL7, CD11c and Ter-119 antibodies and anti-biotin MicroBeads (Miltenyi Biotec, Cat#130-090-485). The enriched splenocytes and BM cells were then sorted for donor-derived PCs (CD45.1^+^SAdump^-^CD138^+^) for scRNA-seq. Memory B-cells were simultaneously sorted as CD45.1^+^SAdump^-^CD138^-^CD19^+^B220^+^CD38^+^FAS^-^IgD^-^ into RPMI (Gibco, cat# 61870). 25,000 MBCs were immediately transferred per sex-matched, CGG-primed wildtype CD45.2^+^ recipient. For CGG priming, mice were injected intraperitoneally with 50 ug of alum-precipitated CGG (Jackson ImmunoResearch, 003-000-002) three weeks prior to MBC transfer. One day after MBC transfer, the recall recipients were injected intraperitoneally with 50 ug of alum-precipitated NP_15_-CGG (Biosearch Technologies, N-5055B) in 100 ul of PBS. Splenic and BM cells (femur and tibia) from the recall recipients were isolated 7 d.p.i. and enriched for donor-derived cells as described above. The enriched splenocytes and BM cells were then sorted for donor-derived PCs (CD45.1^+^SAdump^-^CD138^+^) for scRNA-seq. Chromium Next GEM Single Cell 5’ Gel Bead Kit v2 (10x Genomics) was used for library preparation. Sequencing was performed on the Illumina NovaSeq 6000 and X platforms.

#### scVDJ-seq processing & clonotype calling

Raw VDJ sequencing reads were processed using Cellranger (v7.0.1) and aligned to the refdata-cellranger-vdj-GRCm38-alts-ensembl-7.0.0 and refdata-cellranger-vdj-GRCh38-alts-ensembl-7.1.0 reference genome for mouse and human datasets respectively. The output filtered_contigs.fasta and filtered_contig_annotations.csv files were subsequently reannotated using the Dandelion (v2.0.7) pipeline with Singularity (v3.11.3), where mouse and human BCR sequences were realigned to their respective IMGT reference genome and mutational frequencies were calculated. High-quality BCR contigs were selected by retaining only those with the highest UMI count and at least a two-fold increase in UMI for both heavy and light chains. Clonotypes were identified based on identical VJ gene usage in heavy and light chain contigs across cells with a single pair of heavy and light chain BCR sequences, requiring a minimum of 85% CDR3 amino acid similarity as computed by hamming distance.

#### scRNA-seq processing

FASTQ files of all mouse and human datasets in this study were aligned to the refdata-gex-mm10-2020-A reference genome and refdata-gex-GRCh38-2020-A (Ensembl Release 98) reference genomes respectively, and transcript quantification was performed using the Cellranger pipeline (v7.1.0). Gene expression matrices were processed and analyzed using Seurat (v5.0.1) in R (v4.3.2). For mouse PC datasets, single cells were demultiplexed using the HTODemux() or MULTIseqDemux() functions with default parameters to exclude doublets and empty droplets. For human datasets, demultiplexed outputs from Demuxlet were used to assign cells from individual donors. Genes detected in at least three cells were retained for downstream analysis. Quality control thresholds were applied to exclude cells with fewer than 200 detected genes (nFeature_RNA < 200), high mitochondrial gene content (percent.mt > 10%), detectable haemoglobin transcripts (percent.hb > 0). Additional sample-specific thresholds were applied to remove doublets based on high nCount_RNA and nFeature_RNA (Supplementary Methods). For the mouse timestamped BMPC dataset, cells were removed if the nFeature_RNA and nCount_RNA exceeded 3,000 and 30,000 respectively.

#### scRNA-seq analysis

For each mouse and human dataset, all genes encoding for B-cell receptors (BCRs) were excluded to eliminate the influence of high Immunoglobulin gene expression and isotype usage on cell clustering. Gene expression data were normalized using the SCTransform (v2) method, and the top 3000 highly variable features (HVFs), unless specified otherwise, were selected for sample-level batch correction using Seurat’s RPCA-based integration method. Principal component analysis (PCA) and the Louvain algorithm were employed to identify cell populations initially. Clusters corresponding to contaminating populations were removed: T/NK cells (mouse: Cd3d, Nkg7; human: CD3D, NKG7), B cells (mouse: Cd79a, Ms4a1; human: CD79A, MS4A1), and myeloid cells (mouse: Lyz2, Itgam; human: LYZ, ITGAM). The remaining plasma cell clusters (mouse: Irf4, Prdm1; human: IRF4, PRDM1) were subsequently re-integrated following the same workflow and re-clustered using the Leiden algorithm. For the mouse timestamped BMPC dataset, PC subsets were identified with the top 20 principal components and clustering resolution of 0.3.

#### Label transfer of PC longevity

For all human PC datasets, HGNC symbols were converted to MGI orthologue symbols using gene annotations from Ensembl Release 105. To label plasma cells of varying longevity, we leveraged Seurat’s FindTransferAnchors() and TransferData() functions with the RPCA method. An anchor score threshold of > 0.5 was applied to retain PCs confidently mapped to the reference atlas.

#### Differential expression analysis

Differential gene expression analysis of long-lived plasma cell clusters was carried out using the Seurat’s FindAllMarkers() function, tested with Wilcoxon rank-sum method. Genes were considered significantly differentially expressed based on p_val_adj < 0.05.

#### Gene set enrichment analysis

To determine pathway enrichment, all genes were ranked based on avg_log2FC. Gene set enrichment analysis (GSEA) was performed against MSigDB HALLMARK, KEGG, GO, REACTOME mouse gene set collections using the clusterProfiler package (v4.8.2), with parameters: nperm = 1000, min.overlap = 5.

#### Identification of S2+ PCs

To determine which PCs recognize S2 protein from SARS-CoV-2 virus, we applied a two-step strategy adapted from Lopez de Assis et al. (2023), focusing on BCR sequences of the IGH locus. We first compared V-J usage and CDR3 sequences of each PC to the S2+ MBCs; those V-J combination present in PC pool but absent in the S2+ MBC pool were excluded. Subsequently, we computed all Levenshtein distances of CDR3 amino acid sequence between each remaining PC and the S2+ MBC sharing the same V-J segment. PCs were termed “S2+” if their CDR3 sequence achieved a minimum CDR3 distance of < 0.1 with any S2+ MBCs.

#### Statistical analyses

Statistical methods used in this study are described in figure legends. Comparison was performed using Brown-Forsythe and Welch ANOVA test with Dunnett’s T3 multiple comparisons test using GraphPad Prism10 (GraphPad Software), as well as parametric Welch’s *t*-test under R using ggpubr (v0.6.0). All comparisons are two-tailed unless specified. Sample sizes are indicated in the figures, and significances are shown as exact *P* values or specified in figure legends.

---

**Figure S1.**
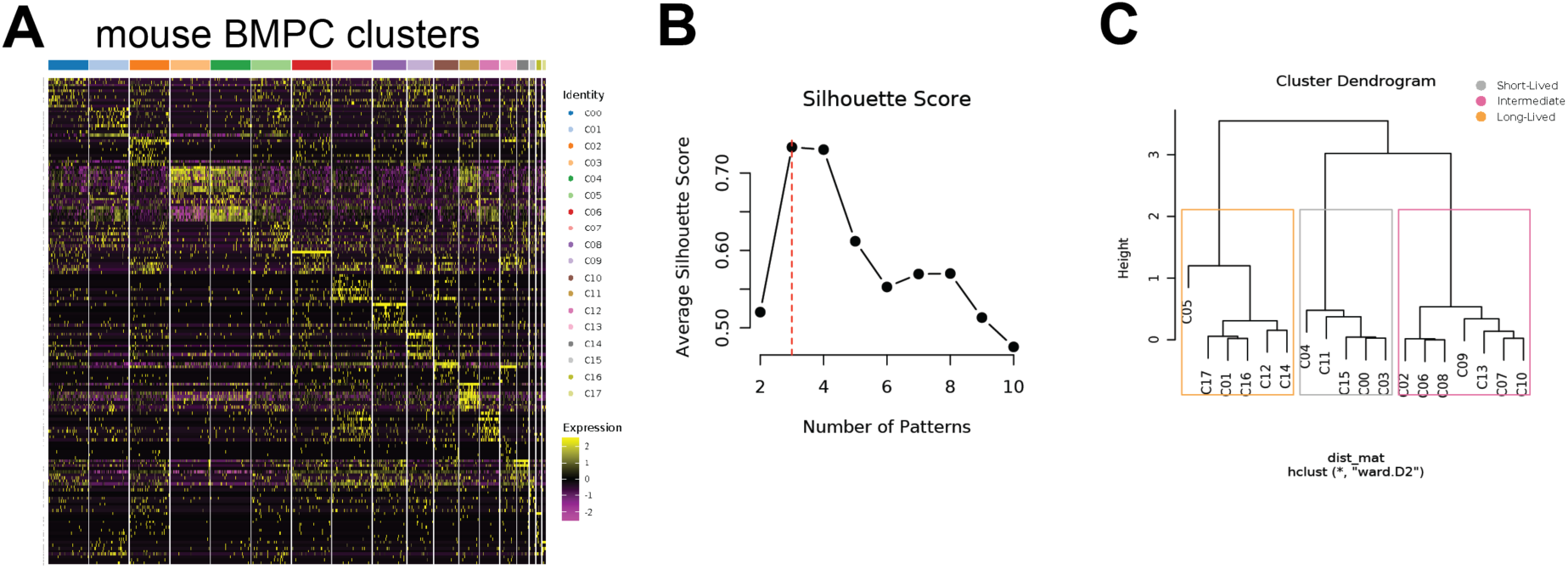
Defining mouse plasma cell longevity states; related to Figure 1. **(A)** Heatmap of top 10 differentially expressed genes per cluster for the bone marrow plasma cell clusters defined in Figure 1. **(B)** Silhouette scores for candidate partitions, used to determine the optimal number of recurrent transcriptional patterns (“longevity programs”). **(C)** Hierarchical clustering dendrogram of clusters based on these patterns, resolving three distinct longevity programs.

**Figure S2.**
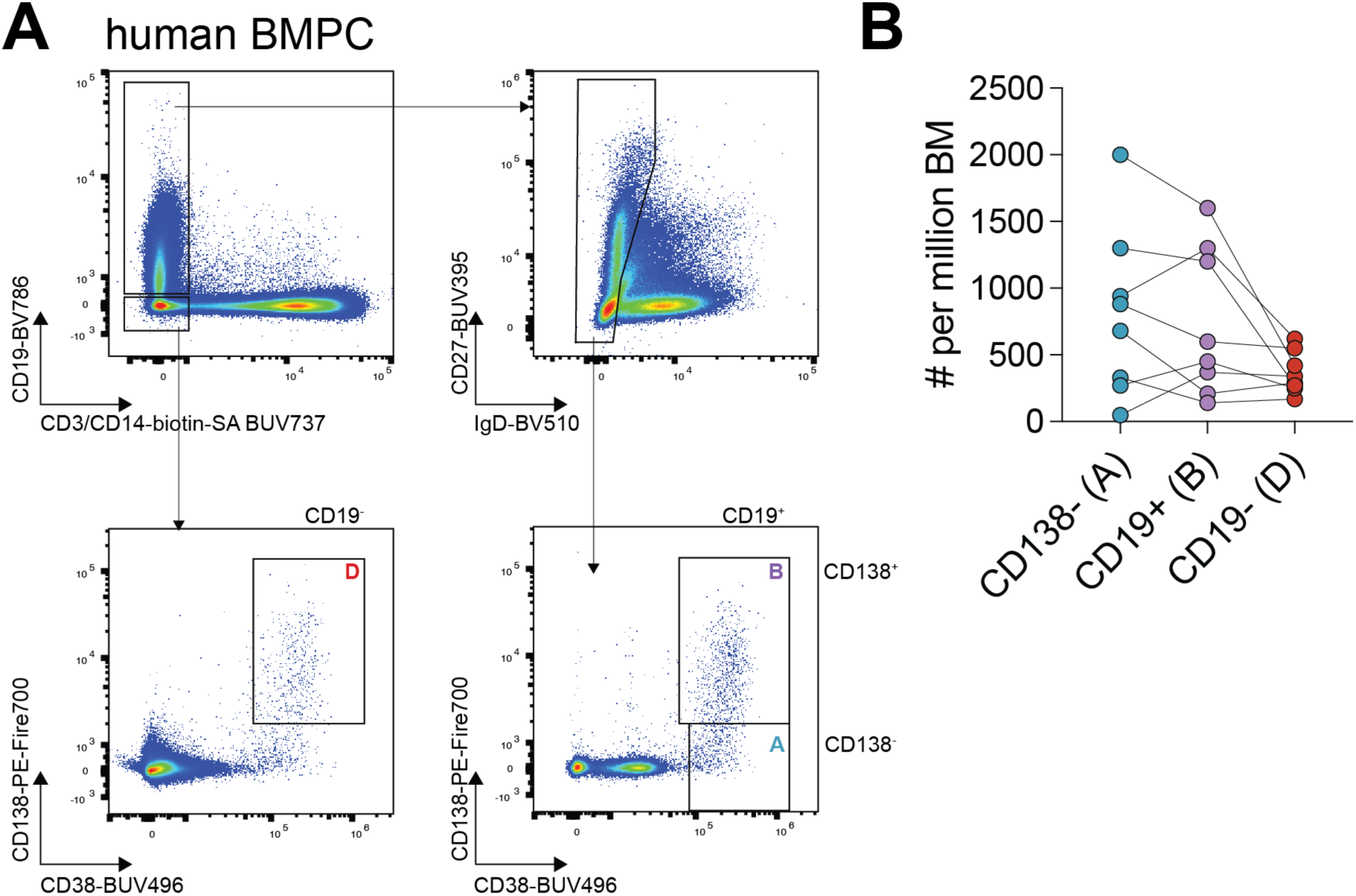
Human bone marrow plasma cell phenotypes; related to Figure 2. **(A)** Flow-cytometric staining and gating strategy used to define human bone marrow plasma cell populations A, B and D. **(B)** Quantification of populations A, B and D, expressed as cell numbers per 10⁶ total human bone marrow cells.

**Figure S3.**
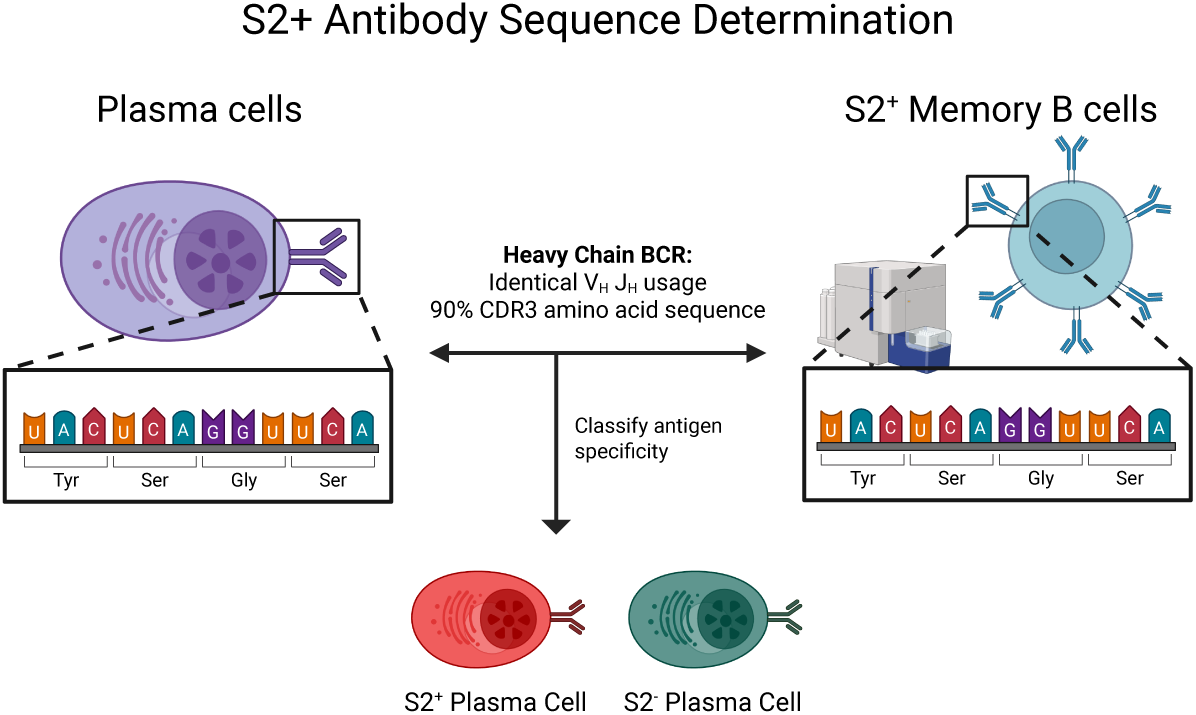
Identification of spike-specific plasma cell clones; related to Figure 4. Schematic illustrating the identification of S2⁺ plasma cells (PCs) by comparing their BCR sequences with those of S2⁺ memory B cells. Created in BioRender. Xu, A., Hung, M. (2025) https://BioRender.com/3be2pso.

**Figure S4.**
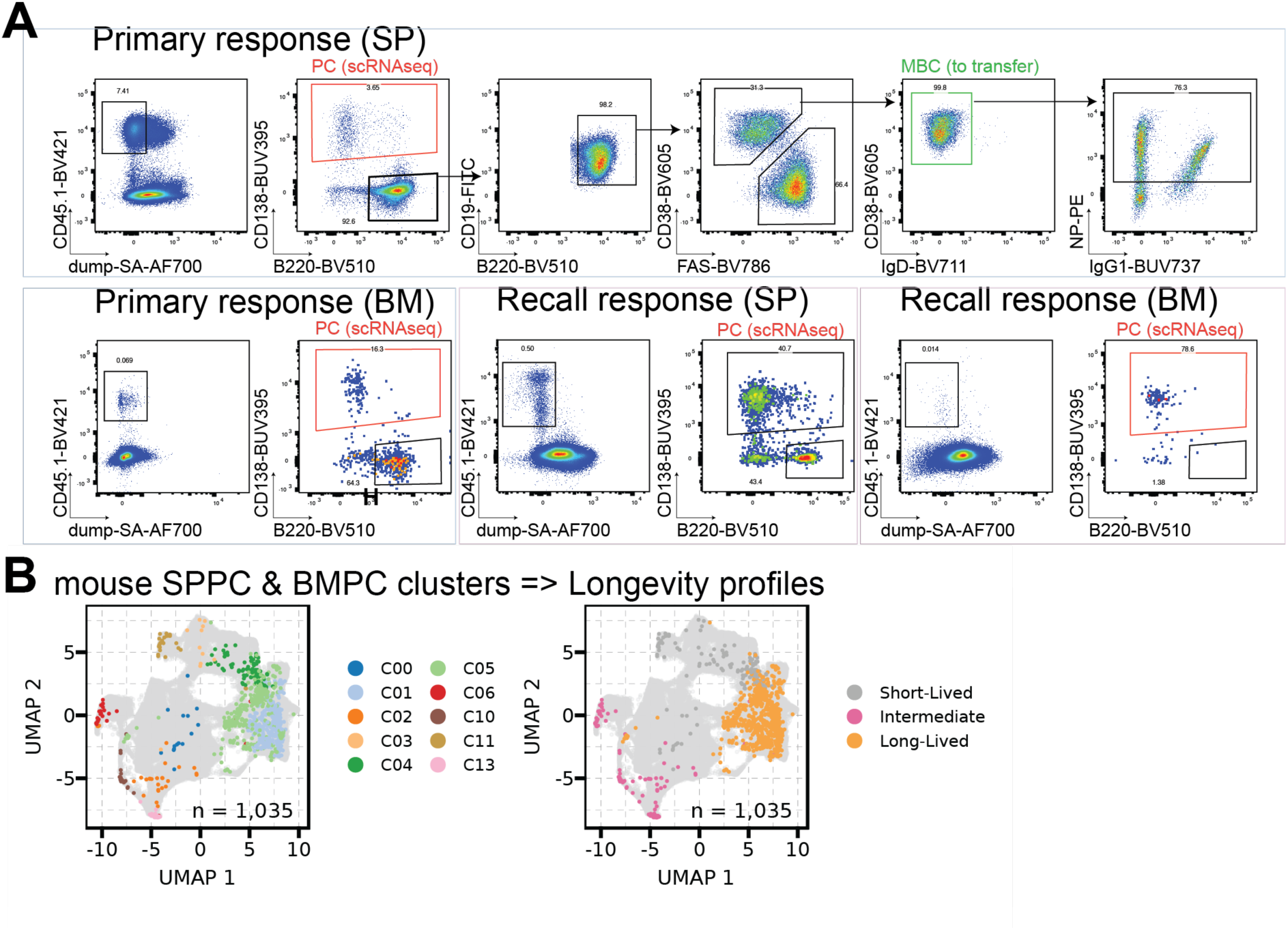
Primary versus MBC-derived plasma cell responses; related to Figure 5. **(A)** Gating strategy for sorting donor-derived plasma cells (PCs) from primary and recall responses for single-cell RNA-seq, and memory B cells (MBCs) from the primary response for adoptive transfer, as shown in Figure 5. **(B)** UMAP projection of splenic (SPPCs) and bone marrow plasma cells (BMPCs) from primary and recall responses, reference-mapped onto longevity clusters and colored by inferred longevity program.

